# Coordinated Assembly of WAVE and WASH complexes

**DOI:** 10.64898/2026.09.18.752629

**Authors:** Nikita M. Novikov, Christine Lazennec-Schurdevin, Peter A. Thomason, Iman Haddad, Lucile Kogey-Fuchs, Magalie Duchateau, Artem I. Fokin, Daria Bondar, Nicolas Dugave, Sergio Lilla, Julia Chamot-Rooke, Joëlle Vinh, Raphaël Guérois, Yves Mechulam, Robert H. Insall, Emmanuelle Schmitt, Alexis M. Gautreau

**Affiliations:** Laboratory of Structural Biology of the Cell (BIOC), CNRS, École Polytechnique, Institut Polytechnique de Paris, 91120 Palaiseau, France; Cancer Research UK Scotland Institute, Garscube Estate, Switchback Road, Glasgow G61 1BD, United Kingdom; Biological Mass Spectrometry and Proteomics (SMBP), ESPCI Paris, PSL University, CNRS UAR2051, 75005 Paris, France; Institut Pasteur, Université Paris Cité, CNRS UAR2024, Mass Spectrometry for Biology, 75015 Paris, France; LCMS Bioanalysis, Laboratory Sciences, Translational Medicine Unit, Sanofi, Vitry-sur-Seine, France; Institute for Integrative Biology of the Cell (I2BC), Université Paris-Saclay, CEA, CNRS, 91198, Gif-sur-Yvette, France; Division of Cell & Developmental Biology, University College London, Biosciences, London WC1E 6BT, United Kingdom

**Keywords:** Arp2/3, NPF, multiprotein complex, cell migration, endosomal trafficking, *Dictyostelium discoideum*

## Abstract

WAVE and WASH polymerize Arp2/3-mediated branched actin at the leading edge of migrating cells and on the surface of endosomes, respectively. These two proteins are regulated within similar multiprotein complexes, the assembly mechanism of which remains poorly understood. Here we found using mass spectrometry that the two smallest subunits, BRK1 of the WAVE complex and CCDC53 of the WASH complex, interact with each other and with the assembly factor HSBP1. HSBP1 promotes WAVE and WASH assemblies and their respective activities, migration persistence and endosomal branched actin. Reduced levels of WAVE and WASH complexes caused by HSBP1 depletion can be corrected by providing cells with an excess of BRK1 or CCDC53 to assemble their specific complex. Unexpectedly, endogenous levels of BRK1 cross-regulate WASH assembly, whereas those of CCDC53 cross-regulate WAVE assembly. We found that various oligomers comprising BRK1, CCDC53 and HSBP1, including a heterotrimer containing one molecule of each, are formed through their promiscuous coiled coils. These oligomers play a coordinating role in the assembly of WAVE and WASH complexes.

## INTRODUCTION

The Arp2/3 complex is a molecular machine that nucleates new actin filaments from the side of pre-existing filaments (Pollard, 2007). This activity results in the formation of dense branched actin networks that produce a pushing force *in vitro* (Loisel *et al*, 1999; Yarar *et al*, 1999). Branched actin networks are found at the proximity of cell membranes and they were shown to generate membrane protrusions from the plasma membrane and to remodel internal membranes (Rotty *et al*, 2013; Molinie & Gautreau, 2018). So-called Nucleation Promoting Factors (NPFs) divide the labor of activating the Arp2/3 at different subcellular locations. For example, WAVE family proteins activate the generation of cortical branched actin in membrane protrusions and are involved in cell migration (Miki *et al*, 1998; Hahne *et al*, 2001; Yamazaki *et al*, 2003), whereas WASH family proteins activate the generation of branched actin at the surface of endosomes and are involved in the scission of transport intermediates containing sorted cargo from endosomes (Derivery *et al*, 2009b; Gomez & Billadeau, 2009).

The two WAVE and WASH NPFs are regulated within stable multiprotein complexes that are distinct, but analogous to one another, with a one-to-one correspondence between their five subunits (Gautreau *et al*, 2004; Derivery *et al*, 2009b; Jia *et al*, 2010). It has been well established that the WAVE NPF is constitutively active, but maintained inactive within its complex (Derivery *et al*, 2009a; Ismail *et al*, 2009; Chen *et al*, 2010, 2017). Most migration signals are thought to activate WAVE through GTP-loading of the RAC1 small GTPase that interacts with the WAVE complex and eventually result in a conformational change exposing the Arp2/3-activating region of the WAVE NPF. For the WASH complex, the model could be similar, but the WASH complex immunopurified from cells keep some NPF activity, because WASH activation involves its non-degradative, K63-mediated poly-ubiquitination (Derivery *et al*, 2009b; Hao *et al*, 2013).

The complexes that regulate WAVE and WASH are critical for the stability of the NPFs. Indeed, the levels of WAVE and WASH are strongly down-regulated when expression of subunits of their respective complexes is targeted by knock-down or knock-out (Kunda *et al*, 2003; Jia *et al*, 2010; Litschko *et al*, 2017). The WAVE NPF is thought to be embedded into its multiprotein complex soon after its translation, because free active WAVE would activate the Arp2/3 throughout the cell and therefore prevent the specific nucleation of branched actin networks, which is finely regulated in time and space (Machesky & Insall, 1998; Derivery *et al*, 2009a). The same reasoning can be applied to WASH. The assembly of WAVE and WASH complexes is therefore thought to be a way for the cell to prevent dominant detrimental effect of free NPFs.

In the cytoplasm, subunits of the WAVE complex are only present in their complexed form, with the notable exception of the smallest subunit BRK1 that also exists in a free form (Gautreau *et al*, 2004). It was demonstrated that free BRK1 is an homotrimer (Derivery *et al*, 2008; Linkner *et al*, 2011), while the BRK1 subunit in the WAVE complex is a single molecule (Chen *et al*, 2010). The BRK1 homotrimer was demonstrated to be a precursor of the assembled BRK1 subunit, indicating that WAVE complex assembly requires the dissociation of the BRK1 homotrimer to contribute a single BRK1 molecule of the WAVE complex (Derivery *et al*, 2008). This transition involves the promiscuity of coiled coils, since the free form of BRK1 is a homotrimeric coiled coil, whereas the WAVE complex implicate a heterotrimeric coiled coil where BRK1, ABI1 and WAVE2 all contribute a single coiled coil strand. Whether this coiled coil transition is spontaneous or require an assembly factor is not known. A single assembly factor has been proposed for the WAVE complex: the Nudel protein, which interacts with BRK1 and other subunits of the WAVE complex (Wu *et al*, 2012), but Nudel implication in this particular transition from homotrimeric to heterotrimeric coiled coils during WAVE complex assembly has not been documented to our knowledge.

For the WASH complex, the only assembly factor identified, HSBP1, interacts with the smallest subunit CCDC53. Both HSBP1 and CCDC53 are organized around trimeric coiled coils (Liu *et al*, 2009; Visweshwaran *et al*, 2018). HSBP1 was shown to facilitate the dissociation of the CCDC53 homotrimer to contribute a single coiled coil strand together with FAM21 and WASH, as initially proposed for the WAVE complex (Derivery *et al*, 2008). Moreover, a previous role attributed to HSBP1 was to inactivate the HSF1 transcription factor, the active form of HSF1 in the nucleus being an homotrimeric coiled coil (Satyal *et al*, 1998). HSBP1 thus appears as a specialized dissociating factor of homotrimeric coiled coils.

Here we identify HSBP1 using proteomics as an assembly factor of the WAVE complex, in addition to its previously established role in assembling the WASH complex. Moreover, we find that CCDC53, BRK1 and HSBP1 form various hetero-oligomeric assemblies around their promiscuous coiled coils. The assembly of both WAVE and WASH complexes is coordinated by these oligomers containing the BRK1 and CCDC53 precursor subunits and the HSBP1 assembly factor.

## RESULTS

### Systematic proteomics of subunits composing WAVE and WASH complexes

To better understand the assembly and mechanisms of action of WAVE and WASH complexes, we looked for partners of their subunits using Tandem Affinity Purification (TAP) and mass spectrometry. To this end, we generated pools of 293 cells stably expressing from the AAVS1 locus N-terminal tagged Flag-GFP fusion proteins of each subunit. The integrative plasmid used drives expression from the strong EF1α promoter. Expression of fusion proteins was verified by Western blots using antibodies targeting GFP or the corresponding subunit. In most cases, expression of the exogenous protein induced a reduction in expression levels of the endogenous protein (Fig.S1). This effect has previously been reported (Derivery *et al*, 2009a) and is interpreted as evidence that the tag does not interfere with WAVE or WASH complex assembly: the excess of the exogenous subunit titrates out endogenous partner subunits in their respective complex and unincorporated subunits are then probably degraded (Derivery & Gautreau, 2010; Pla-Prats & Thomä, 2022). The only exception was BRK1, which also exists as a free pool (Gautreau *et al*, 2004).

We then performed TAP using Flag immunoprecipitation, then eluting Flag beads with Flag peptides and finally immunoprecipitating the dually tagged subunits using GFP trap beads, i.e. beads covalently coupled to a monomeric GFP binding protein. These two sequential immunoprecipitations reduce contaminants compared with a single immunoprecipitation. The bait and its partners released from GFP trap beads were examined by SDS-PAGE (Fig.1A). The rest of the immunoprecipitate was analyzed in bulk by mass spectrometry using LC-MS/MS and Label-Free Quantification (LFQ). For each subunit of WAVE and WASH complexes, hundreds of potential partners were identified (Table S1; Table S2). In sharp contrast, only a few occasional peptides were retrieved from cells expressing FlagGFP.

**Figure 1.**
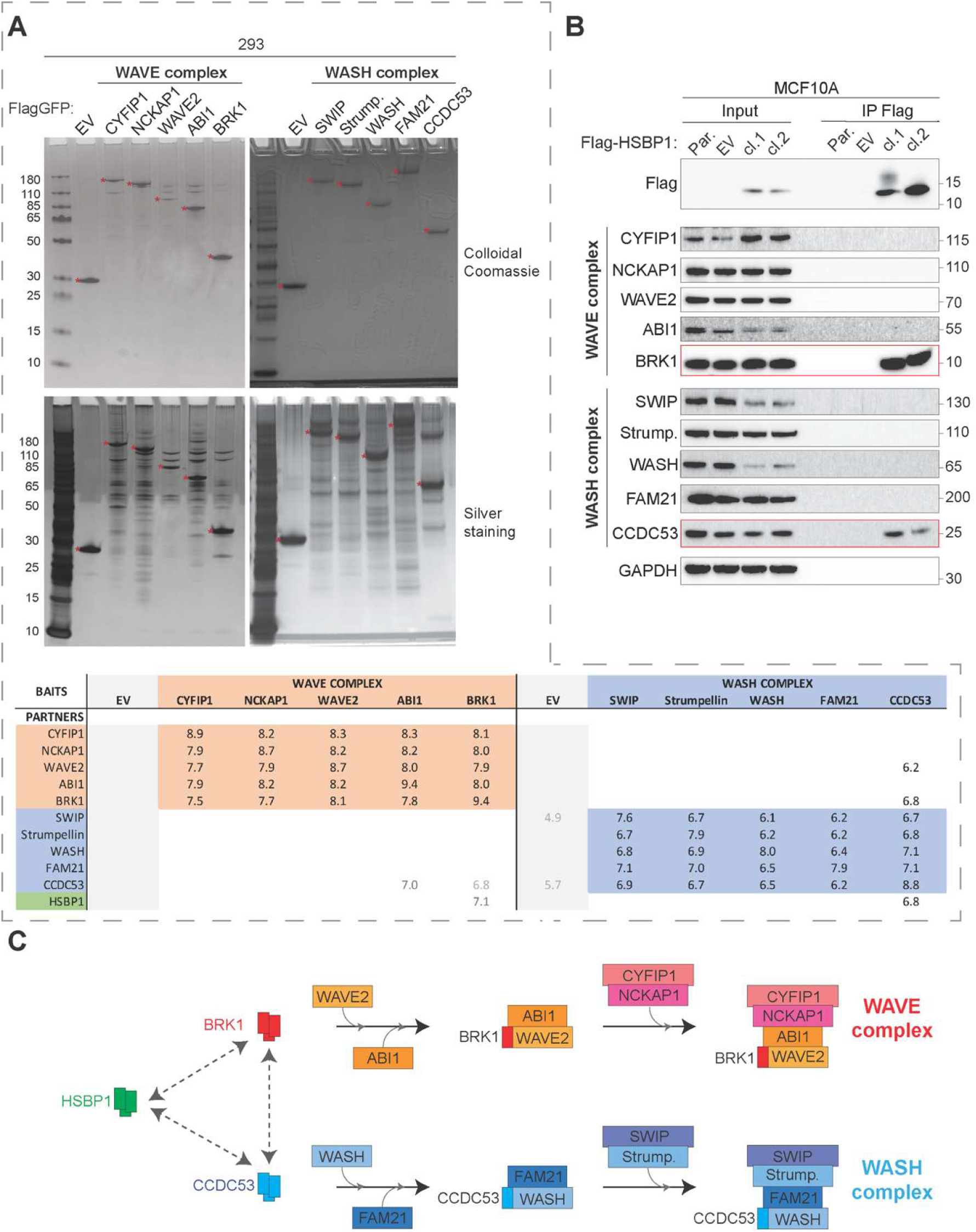
Proteomics of the subunits of WAVE and WASH complexes. **(A)** Stable 293 cell lines expressing Flag-GFP or Flag-GFP tagged subunits of WAVE or WASH complexes are subjected to Tandem Affinity Purification (TAP). Partner proteins are resolved by SDS–PAGE and revealed using colloidal Coomassie or silver staining. Partner proteins are identified and quantified by mass spectrometry using LC-MS/MS and Label-Free Quantification (LFQ). Red dots indicate the position of bait proteins. Table recapitulating quantification of interacting partners (average of Log10 of LFQ intensity in three independent biological repeats). Numbers in black correspond to proteins identified in three repeats out of three; intermediate gray in two repeats only; light gray in a single repeat. **(B)** Lysates from two independent clones expressing Flag-tagged HSBP1 and controls are subjected to Flag immunoprecipitations. Lysates and immunoprecipitates are analyzed by Western blot using the indicated antibodies. Par.: Parental cells. EV: Empty Vector. **(C)** Scheme representing the parallel roles of HSBP1 in assembling the analogous WAVE and WASH complexes through tripartite interactions between BRK1, CCDC53 and HSBP1.

We first verified that each WAVE complex subunit qualitatively retrieved all other subunits of the WAVE complex and that each WASH complex subunit qualitatively retrieved all other subunits of the WASH complex (Fig.1A). These observations confirm that the fusion of subunits with FlagGFP does not prevent the assembly of their respective complex. The overexpressed bait protein was systematically more abundant in the TAP than any other partner subunits, suggesting that in this cell system, a significant fraction of the bait is not incorporated into its complex.

The amount of WAVE complex retrieved by one of its tagged subunits, estimated by the LFQ of partner subunits, was systematically larger than the amount of WASH complex retrieved by a tagged subunit, indicating that the WAVE complex is more abundant in 293 cells than the WASH complex (Fig.1A). They were a couple of unexpected interactions, of relatively low abundance, specifically between CCDC53 and WAVE complex subunits. The TAP of CCDC53 systematically retrieved WAVE2 in three out of three biological repeats, but the reciprocal TAP was not functioning at all: CCDC53 was not found in the three TAP experiments of WAVE2. Similarly, the TAP of ABI1 retrieved CCDC53 in two repeats out of three, but here again, the reciprocal TAP did not function. Finally, the TAP of CCDC53 retrieved BRK1 in three out of three biological repeats. This time, the reverse TAP of BRK1 retrieved CCDC53 in one of the three biological repeats.

An expected interaction of CCDC53 was with HSBP1, which our group had previously identified as an assembly factor for the WASH complex (Visweshwaran *et al*, 2018). HSBP1 interacts with CCDC53, but with no other subunit of the WASH complex, indicating that it binds to CCDC53 before the WASH complex is assembled, not afterwards. We also identified HSBP1 as a specific partner of BRK1, but to no other subunit of the WAVE complex (Fig.1A). To validate these interactions, we isolated two stable MCF10A clones that express Flag-HSBP1 to perform the reverse immunoprecipitation. Indeed, Flag immunoprecipitation confirmed that HSBP1 interacts with both CCDC53 and BRK1, but not with any other subunits of the WASH and WAVE complexes (Fig.1B). This result suggests that HSBP1 may participate in the assembly of the WAVE complex, in addition to its already established role in the assembly of the WASH complex (Fig.1C). The interaction of HSBP1 with both BRK1 and CCDC53 can also bridge the two proteins and thereby potentially account for the CCDC53-BRK1 interaction that we have detected here.

### HSBP1 promotes WAVE and WASH assemblies and their respective activity

To investigate the role of HSBP1, we generated HSBP1 knockout (KO) clones in MCF10A cells using CRISPR/Cas9. HSBP1 is a small protein of only 76 amino-acids and its detection by Western blot is difficult. Even if a specific blotting procedure has been developed (Visweshwaran *et al*, 2018), detection of the endogenous HSBP1 protein remains at the limit, rendering its use as a screening method risky. We therefore used the transfection of two gRNAs cutting on both sides of the starting ATG codon and a third gRNA that targets *ATP1A1* as a way to select genome-edited cells with ouabain (Agudelo *et al*, 2017). This procedure allows efficient selection of clones harboring deletions in MCF10A cells (Fokin *et al*, 2025). Two independent KO clones were identified using PCR on genomic DNA and confirmed by sequencing (Fig.S2). As expected, we did not detect HSBP1 expression in these two clones by Western blot, contrary to parental cells and control cells that were selected to resist ouabain (Fig.2A). As previously reported, HSBP1 inactivation induced a down-regulation of WASH and CCDC53 (Visweshwaran *et al*, 2018), but also down-regulated WAVE complex subunits (Fig.2A): HSBP1 KO induced a down-regulation of WAVE2 and BRK1, the subunits of the WAVE complex that correspond to WASH and CCDC53 in the WASH complex, as well as the remaining subunits, CYFIP1, NCKAP1 and ABI1, of the WAVE complex.

**Figure 2.**
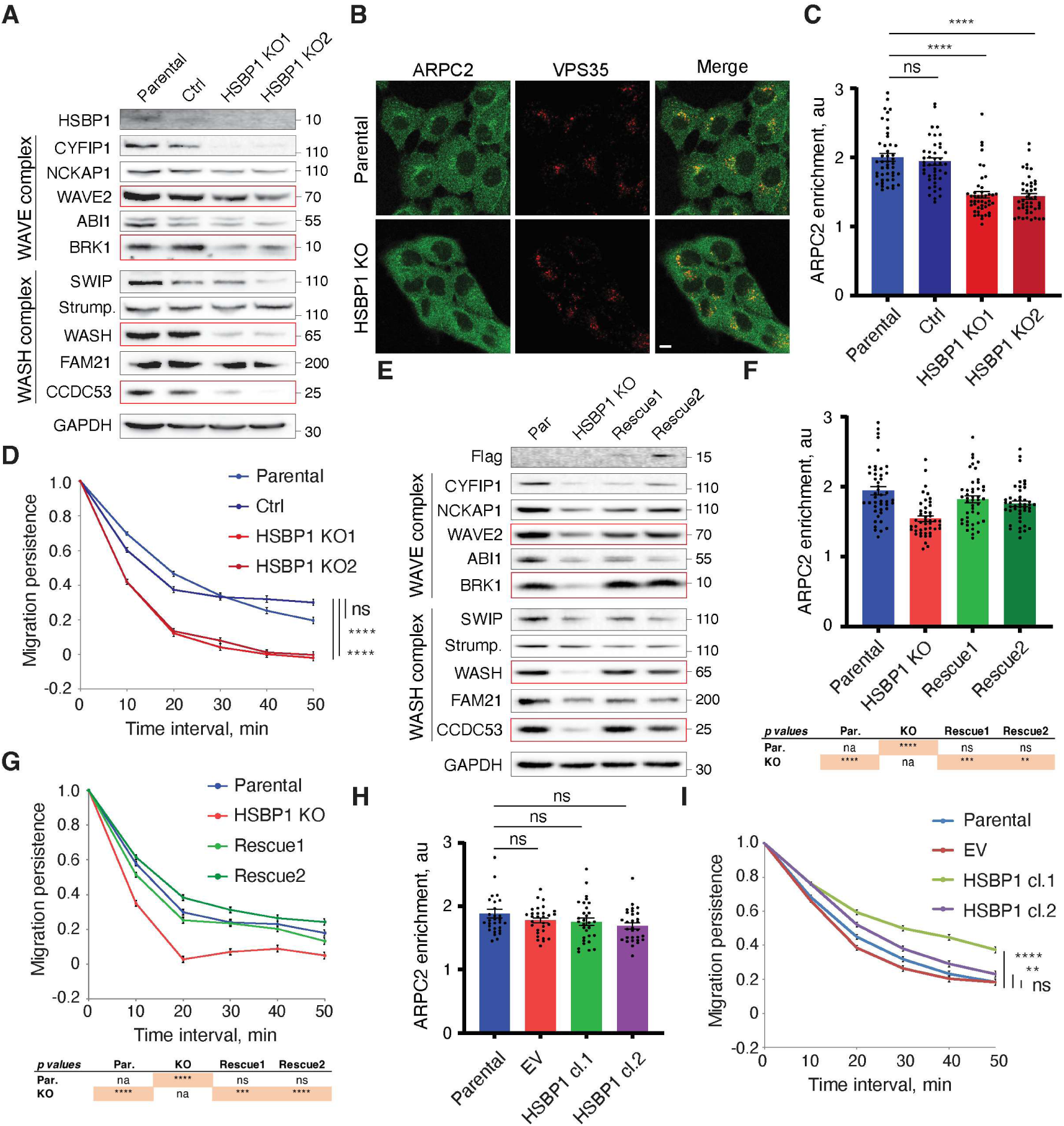
HSBP1 promotes the activities that depend on the WAVE and WASH. **(A)** MCF10A parental cells, control cells and two HSBP1 KO clones are analyzed for the levels of WAVE and WASH complex subunits by Western blots using indicated antibodies. **(B)** HSBP1 KO cells are stained with antibodies against ARPC2 and VPS35 by immunofluorescence. The ARPC2 subunit of the Arp2/3 complex reveals WASH-dependent branched actin networks at the surface of endosomes that are positive for the retromer subunit VPS35. Single confocal section, scale bar 10 µm. **(C)** Quantification of endosomal ARPC2 enrichment in HSBP1 KO cells. Three biological repeats with similar results. Mean ± SEM, n=45 cells per condition; Kruskal-Wallis followed by Dunn’s multiple comparison tests. **(D)** Migration persistence of single HSBP1 KO cells. Three biological repeats with similar results. Mean ± SEM, n=60 cells. Statistical significance of persistence is analyzed in a non-linear mixed effect model. **(E)** MCF10A parental cells, HSBP1 KO and two rescued clones, i.e. HSBP1 KO cells stably expressing Flag-HSBP1 are analyzed by Western blots using indicated antibodies. **(F)** Quantification of ARPC2 enrichment on the surface of endosomes in rescued clones. Three biological repeats with similar results. Mean ± SEM, n=45 cells per condition; Kruskal-Wallis followed by Dunn’s multiple comparison tests, table reporting the p-values. **(G)** Migration persistence of single rescued cells. Three biological repeats with similar results. Mean ± SEM, n=60 cells. Statistical significance of persistence is analyzed in a non-linear mixed effect model. Table reporting the p-values. **(H)** MCF10A parental cells, two stable clones overexpressing Flag-HSBP1, or the empty vector (EV) as a control, are stained with antibodies against ARPC2 and VPS35 by immunofluorescence. Quantification of ARPC2 enrichment on the surface of endosomes in overexpressing clones. Three biological repeats with similar results. Mean ± SEM, n=30 cells per condition; Kruskal-Wallis followed by Dunn’s multiple comparison tests. Table reporting the p-values. **(I)** Migration persistence of single rescued cells. Three biological repeats with similar results. Mean ± SEM, n=60 cells. Statistical significance of persistence is analyzed in a non-linear mixed effect model. ** p<0.01, *** p<0.0001, **** p<0.0001, ns non-significant, na not applicable.

We next examined the functional consequences of HSBP1 loss on WAVE- and WASH-dependent cellular processes. WASH recruits and activates the Arp2/3 complex at the surface of endosomes at the level of microdomains enriched in the retromer complex (Derivery *et al*, 2009b; Gomez & Billadeau, 2009; Jia *et al*, 2012; Helfer *et al*, 2013). In contrast, the Arp2/3 complex activated by WAVE is critical for membrane protrusions and cell migration. In single MCF10A cells, the major migration parameter that this pathway controls is persistence (Molinie *et al*, 2019). In line with the idea that HSBP1 promotes the assembly of both WASH and WAVE complexes, HSBP1 KO clones were significantly impaired in Arp2/3 recruitment at the surface of endosomes (Fig.2B,C) and in migration persistence (Fig.2D, Fig.S3, Movie S1).

We then expressed Flag-tagged HSBP1 in HSBP1 KO cells and established two independent rescue clones. Re-expression of HSBP1 restored the levels of the NPFs and the small subunits for both WAVE and WASH complexes, BRK1 and WAVE2, or CCDC53 and WASH, respectively (Fig.2E). Rescue on levels of CYFIP1, NCKAP1 and ABI1 in HSBP1 KO cells was not as complete as that of BRK1 and WAVE2. Functionally, HSBP1 re-expression restored branched actin at the surface of endosomes (Fig.2F), and migration persistence (Fig.2G and Fig.S3). Overexpression of HSBP1 in parental MCF10A cells did not enhance endosomal branched actin (Fig.2H), but increased migration persistence compared with parental cells, or cells transfected with the empty vector (Fig.2I). Migration persistence is therefore a cell parameter more sensitive to HSBP1 dosage than the amount of endosomal branched actin.

Finally, we asked whether the increase of subunit levels by HSBP1 could be compensated by the direct expression of the small subunits of either complex. To test this, we stably expressed FlagGFP-BRK1 or FlagGFP-CCDC53 in MCF10A HSBP1 KO cells and analyzed levels of the two complexes by Western blot. First we noticed that the overexpression of tagged CCDC53 drastically decreased levels of endogenous untagged CCDC53 (Fig.3A,B). The same effect was observed for BRK1, albeit not as pronounced. This replacement of endogenous subunits by tagged subunits was already shown when clones expressing these subunits were isolated for TAP in parental cells (Fig.S1). The most important effect, however, was at the level of the NPF itself: overexpression of CCDC53 selectively restored WASH levels and BRK1 restored WAVE levels to the levels displayed in parental cells (Fig.3A,B). The effects of BRK1 and CCDC53 overexpression on the other subunits of the two complexes were not statistically significant (Fig.S4). This experiment reveals that HSBP1 is not absolutely required for the assembly of either the WAVE or the WASH complex, but that it facilitates both (Fig.3C).

**Figure 3.**
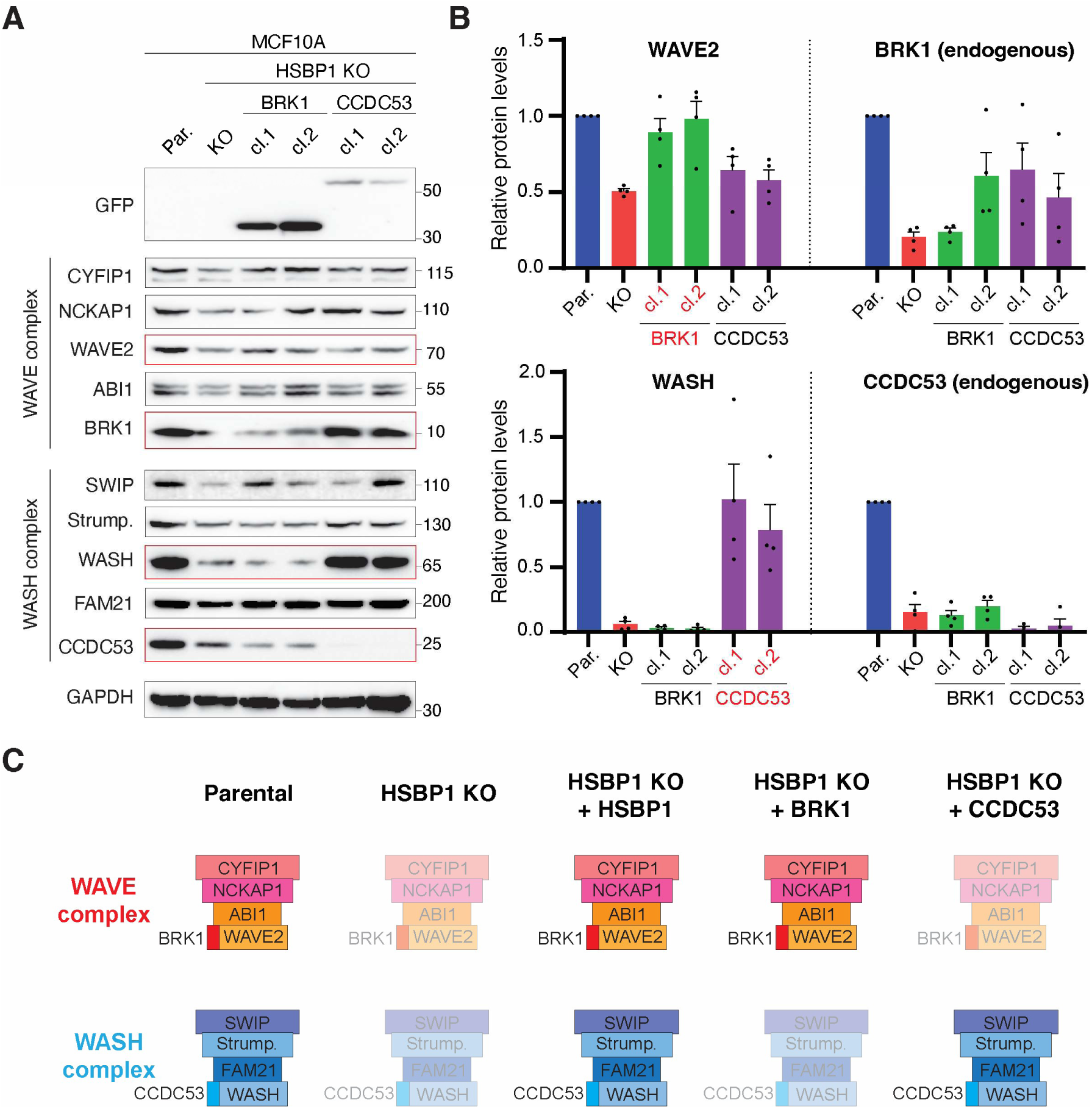
In the absence of HSBP1, overexpression of BRK1 or CCDC53 rescues levels of WAVE and WASH complexes, respectively. **(A)** MCF10A parental cells, HSBP1 KO cells and two clones of each HSBP1 KO cells overexpressing either BRK1 or CCDC53 are analyzed by Western blots with indicated antibodies. **(B)** Quantification of Western blots by densitometry, for selected proteins. Four biological repeats, mean ± SEM. **(C)** Levels of WAVE and WASH complexes, based on the amount of the NPF, are illustrated by transparency of the schemes.

We previously reported that the function of HSBP1 as an assembly factor of the WASH complex was conserved in the amoeba *Dictyostelium discoideum* (Visweshwaran *et al*, 2018). We sought to check whether HSBP1 was also an assembly factor for the SCAR complex in the amoeba. SCAR is the homolog of WAVE in the amoeba (Bear *et al*, 1998). One of the phenotypes of the SCAR KO is to decrease the speed of cell migration (Blagg *et al*, 2003). We therefore checked the speed of wild type and HSBP1 KO amoebas in a chemotaxis assay. Cell speed was indeed decreased in the HSBP1 KO amoeba compared with the parental strain (Fig.4A, Movie S2), as if SCAR was depleted. We therefore compared the subunit levels of both SCAR and WASH complexes. In the absence of antibodies recognizing these multiple proteins in the amoeba, we turned to mass spectrometry to identify differentially expressed proteins. To this end, we alternatively dimethyl labeled peptides from one condition (WT or HSBP1 KO) and compared this condition to the other by mixing equal amounts of peptides. This approach revealed a dozen differentially expressed proteins (Table S3). WASH1, SCAR and BRK1 was among them. CCDC53 was, unfortunately, not detected in this experiment. The two NPFs SCAR and WASH were profoundly down-regulated in HSBP1 KO amoebae, less than 10 % of the levels displayed by wild type amoebae (Fig.4B). As in human cells, the other subunits were less depleted in HSBP1 KO amoebas than the NPFs themselves. We validated the depletion of SCAR by Western blot and expression of HSBP1-GFP in the HSBP1 KO amoeba was found to rescue SCAR levels (Fig.4C). The expression of BRK1-GFP in the KO amoeba also rescued expression of SCAR (Fig.4D), as in human cells. We concluded that the function of HSBP1 in promoting the assembly of SCAR/WAVE complexes is evolutionarily conserved. This conserved role of HSBP1 is, however, not absolutely required, since it can be circumvented in amoebae as in human cells by the overexpression of the small subunit BRK1.

**Figure 4.**
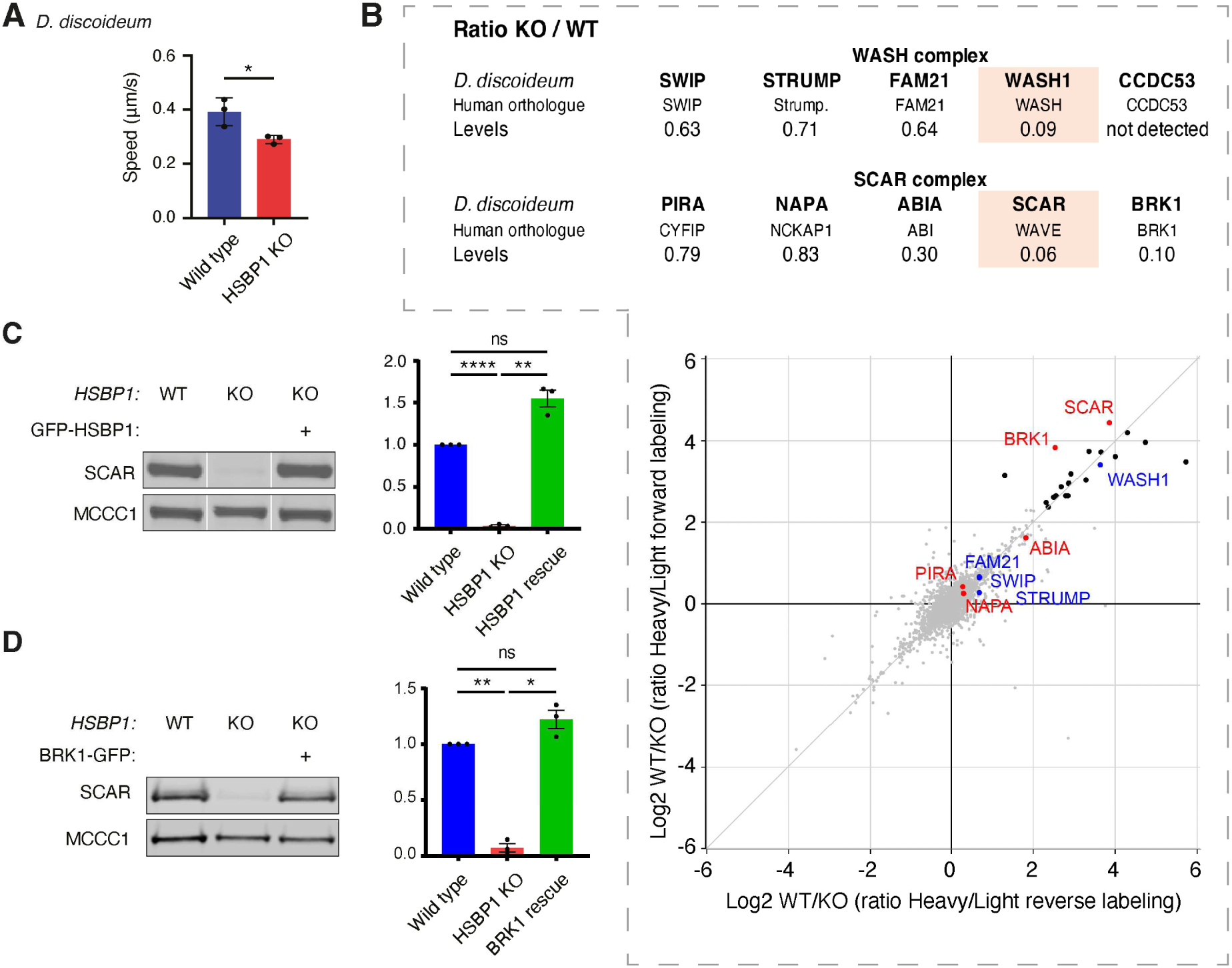
HSBP1 promotes assembly of the SCAR/WAVE complex in *Dictyostelium discoideum*. **(A)** Amoebae are tested in an under agar chemotactic assay using folate as a chemoattractant. Average instantaneous speed of cells. Three biological repeats with similar results, Student’s t-test. **(B)** Lysates prepared from either parental or HSBP1 KO amoeba are alternatively dimethyl-labeled and compared by mass spectrometry to identify differentially expressed proteins. SCAR/WAVE and WASH are depleted from HSBP1 KO amoebae compared with parental ones. **(C)** Parental, HSBP1 KO, and HSBP1 rescued amoebas are analyzed by Western blots for levels of the SCAR/WAVE NPF and MCCC1 as a loading control. Quantification by densitometry mean ± SEM, three biological repeats. ANOVA/Tukey. **(D)** Rescue of SCAR/WAVE levels upon expression of BRK1-GFP. * p<0.05, ** p<0.01, **** p<0.0001, ns non-significant.

### BRK1, CCDC53 and HSBP1 assemble various hetero-oligomers

BRK1, CCDC53 and HSBP1 are three small proteins (75, 194, 76 amino-acids, respectively). They have all been described being organized around coiled coils that preferentially form homotrimers (Tai *et al*, 2002; Derivery *et al*, 2008; Chen *et al*, 2010; Jia *et al*, 2010; Linkner *et al*, 2011; Visweshwaran *et al*, 2018). Coiled coils are often promiscuous and we have already reported that combining purified CCDC53 with HSBP1 spontaneously produces mixed heterotrimers (Visweshwaran *et al*, 2018). Given the association of these three proteins in TAP experiments, we first examined whether their coexpression allowed the formation of trimers containing each protein (see Supplementary Methods). Upon co-expression of His tagged HSBP1 with PC-BRK1 and Flag-CCDC53, the retrieval of His-HSBP1 by pull-down retrieved BRK1, but not CCDC53 (Fig.S5). When the pull-down was further resolved by gel filtration, two peaks containing both HSBP1 and BRK1 were observed, indicating that the pull-down contains more than one molecular species.

To investigate the nature of the oligomers present in this experiment, we turned to native mass spectrometry. The analysis of the major peak after gel filtration (peak 1) provided a spectrum, where a heterodimer composed of one molecule of His-HSBP1 and one molecule of PC-BRK1 was unambiguously identified (Fig.5A). The analysis of peak 2 by native mass spectrometry did not identify any hetero-oligomer. To reconstitute complexes containing CCDC53, we purified individual recombinant CCDC53, BRK1 and HSBP1 proteins from *E. coli* and mixed them after purification. We mixed and incubated HisFlag-CCDC53 with a two-fold molar excess of either PC tagged BRK1 or untagged HSBP1, or both. We characterized by native mass spectrometry the material pulled down by metal beads through the His-tagged CCDC53 (Fig.S6). When the three proteins were mixed, we identified two trimers: one heterotrimer containing one molecule of each, and one containing two PC tagged HSBP1 and one HisFlag-CCDC53. A dimer of HisFlag-CCDC53 was also observed. When HisFlag-CCDC53 was mixed with untagged HSBP1, we identified the two possible heterotrimers, in a one to two ratio or a two to one ratio (Fig.5A). Finally, when HisFlag-CCDC53 was mixed with PC-BRK1, we detected a dimer containing one molecule of each. A dimer of PC-BRK1 was also identified. For both individual proteins and protein complexes, a good match between measured and theoretical masses is observed. Our analyses showed that BRK1 forms a covalent dimer both in native and denaturing conditions. The formation of disulfide bridges between BRK1 and CCDC53 cannot be excluded.

**Figure 5.**
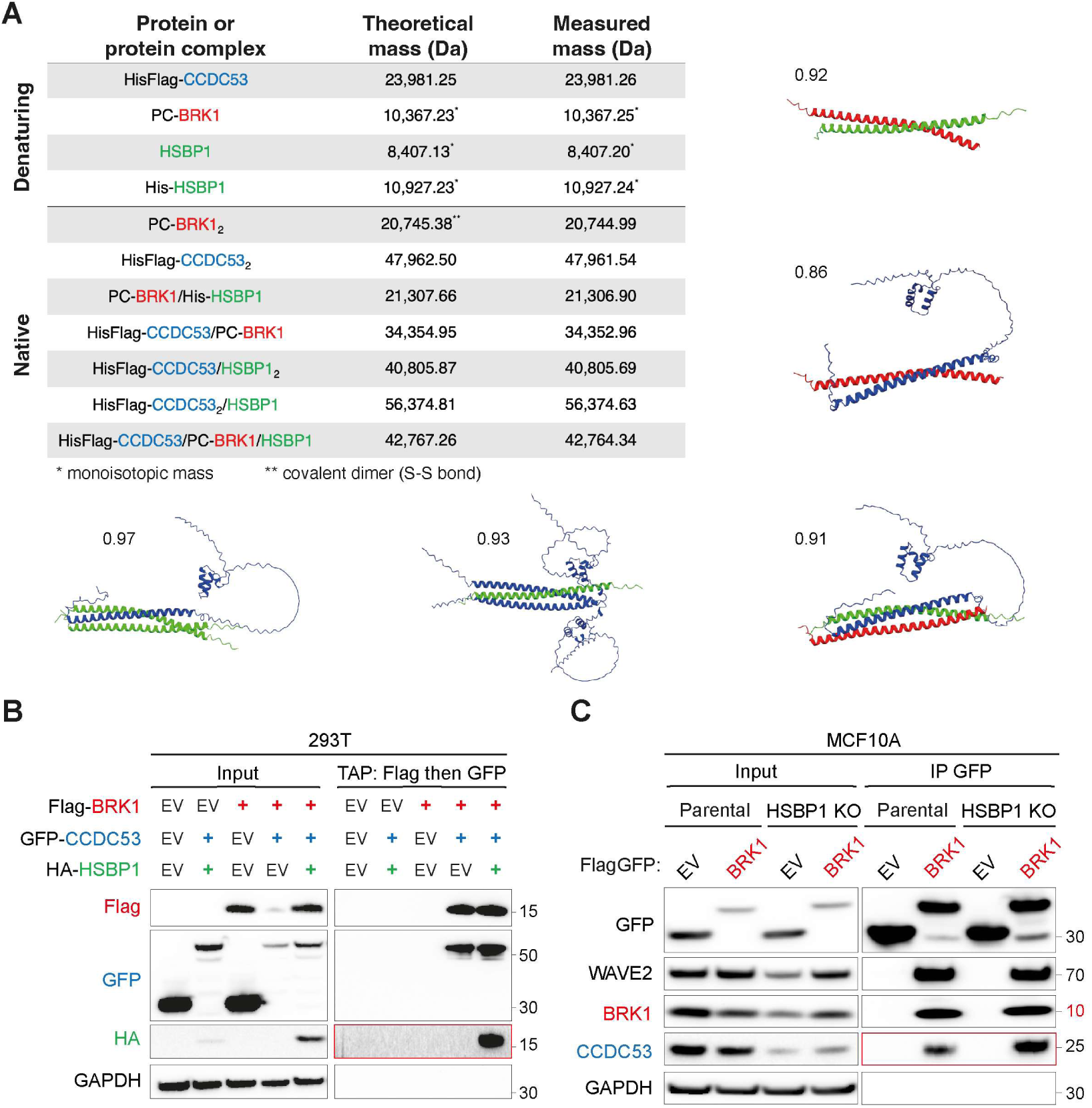
Flexibility in the assembly of HSBP1/BRK1/CCDC53 hetero-oligomers. **(A)** Recombinant BRK1, CCDC53 and HSBP1 proteins are purified from *Escherichia coli.* Full-length proteins and reconstituted complexes are characterized by mass spectrometry in denaturing or native conditions. Five hetero-oligomers, including the HSBP1/BRK1/CCDC53 heterotrimer, are detected by native mass spectrometry and structurally modeled using AlphaFold2. The confidence of the predicted interactions is assessed using actifpTM values, indicated next to each model. **(B)** Existence of the HSBP1/BRK1/CCDC53 heterotrimer in the cell. 293T cells are transiently co-transfected with plasmids expressing Flag-BRK1, GFP-CCDC53 and HA-HSBP1, or their respective control plasmids with no insert (EV: Empty Vector), allowed to grow for 48 hours and then subjected to Flag-GFP tandem affinity purification (TAP) and analyzed by Western blots using indicated antibodies. **(C)** BRK1 can hetero-oligomerize with CCDC53 without HSBP1. The MCF10A parental line and a MCF10A HSBP1 KO clone that stably express FlagGFP-BRK1 at similar levels are subjected to GFP immunoprecipitation. Western blots using the indicated antibodies. Three biological repeats of experiments displayed in B and C were performed and yielded similar results.

Models of all detected hetero-oligomers were obtained using AlphaFold2. These models of hetero-dimers or hetero-trimers showed a critical role of coiled coils, similar to that observed in the structures of individual proteins. Using actifpTM, the metric of the actual interface pTM (Varga *et al*, 2025), that is appropriate for coiled coils and proteins with disordered regions like CCDC53, all models showed high confidence (Fig.5A; actifpTM >0.85).

The heterotrimer containing CCDC53, BRK1 and HSBP1 we detected in vitro is an attractive structure to explain that all three of these proteins binds to the two others and that HSBP1 might dissociate trimers of CCDC53 and BRK1 to provide a single molecule of the small subunit for the assembly of WAVE and WASH complexes. We thus sought to confirm that such a heterotrimer exists in the cell. To this end, we coexpressed in 293T cells, Flag-BRK1, GFP-CCDC53 and HA-HSBP1, and submitted lysates from transfected cells to TAP using sequential Flag and GFP immunoprecipitations (Fig.5B). TAP selected hetero-oligomers containing Flag-BRK1 and GFP-CCDC53. The third protein, HA-HSBP1 was present in the TAP fraction, indicating that at least some of the oligomers selected by the TAP can accommodate the third protein, HSBP1, in the same structure, most likely the hetero-trimer observed using native mass spectrometry of reconstituted complexes.

We also sought to verify in cells that BRK1 can form oligomers with CCDC53, even when HSBP1 is not present, in line with some of the reconstituted complexes characterized using native mass spectrometry, but also to account for the assembly of WAVE and WASH complexes in the absence of HSBP1. To this end, we expressed FlagGFP-BRK1 or the empty vector as a control, in one of the HSBP1 KO MCF10A clones. The absence of HSBP1 did not prevent the formation of BRK1 / CCDC53 oligomers, nor the incorporation of FlagGFP-BRK1 into the WAVE complex (Fig.5C). Together, reconstitution experiments of hetero-oligomers using recombinant proteins and expressions of tagged proteins in cells indicate that a variety of oligomers can be formed between BRK1, CCDC53 and HSBP1 with an apparent versatility in composition and stoichiometry.

### Cross-regulation of WAVE and WASH assembly by the small subunit of the other complex

We reasoned that if mixed oligomers of BRK1, CCDC53 and HSBP1 are the forms that provide the precursor subunits for WAVE and WASH complex assembly, then WAVE complex assembly might depend on CCDC53 levels, and, reciprocally, WASH complex assembly might depend on BRK1 levels. We therefore depleted MCF10A cells of either BRK1, CCDC53 or HSBP1, using pools of siRNAs, and monitored levels of both complexes using Western blots and densitometry in four biological replicates (Fig.6A, Fig.S7). For each NPF, the most dramatic effect on complex assembly was observed when the cognate small subunit was depleted: WAVE levels were most decreased upon BRK1 depletion and WASH levels were most decreased upon CCDC53 depletion. siRNA-mediated depletion of HSBP1 that affected both WAVE and WASH complexes was second in the amplitude of the effect. HSBP1 depletion affected the levels of all subunits of the WAVE complex, but only WASH and CCDC53 of the WASH complex, in line with results obtained upon HSBP1 KO. A third order effect was significant: the effect of the small subunit of one complex on the assembly of the other. We found that CCDC53 depletion significantly decreased the levels of WAVE complexes, whereas that of BRK1 significantly decreased the levels of WASH complexes (Fig.6B), revealing an unexpected cross-talk between the assembly of the two complexes.

**Figure 6.**
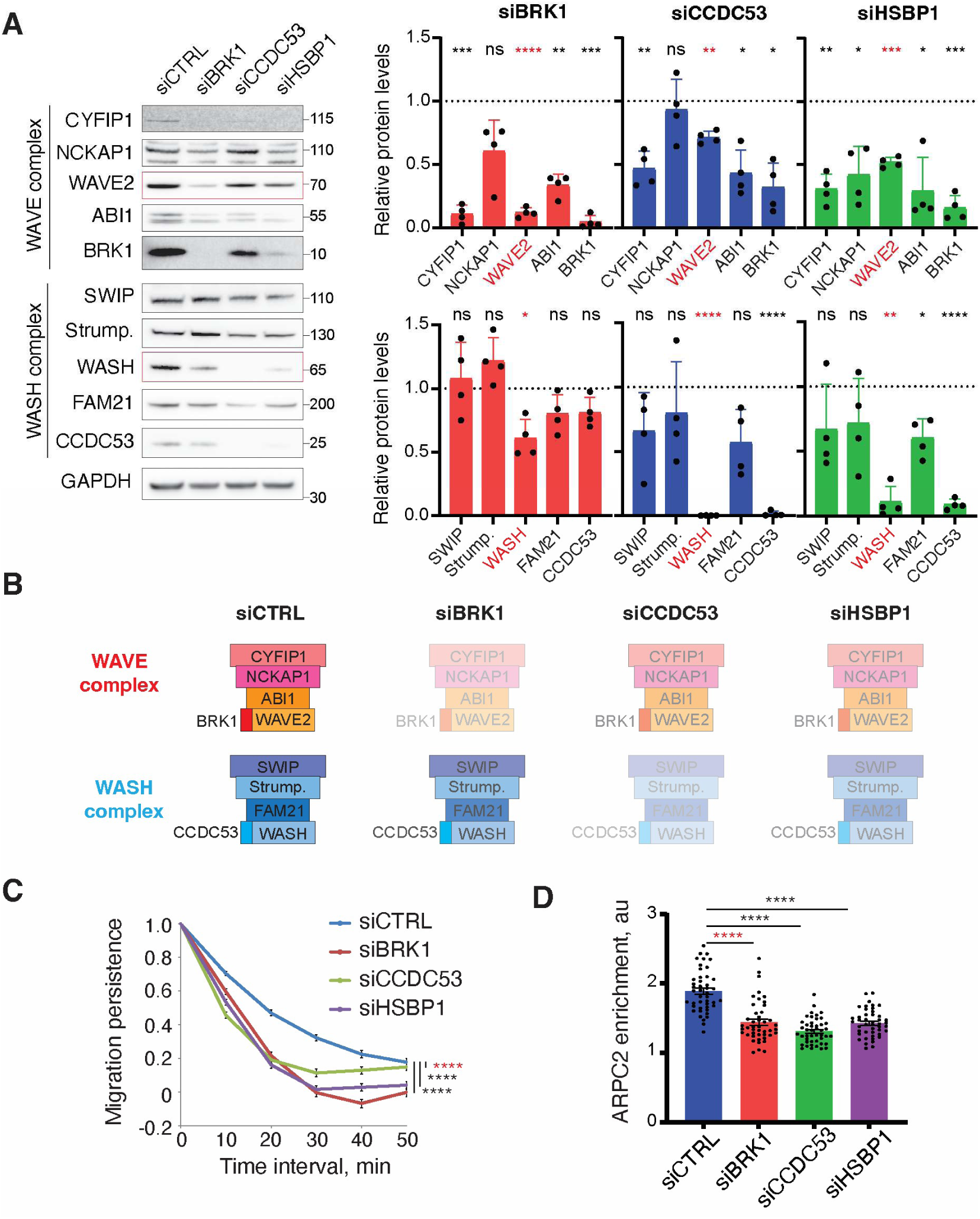
BRK1 cross-regulates WASH activity and CCDC53 cross-regulates WAVE activity. MCF10A cells are transfected with pools of siRNAs targeting BRK1, CCDC53 or HSBP1, or non-targeting siRNAs as a control, and analyzed 3 days after transfection. **(A)** Cell lysates are analyzed by Western blots. Quantification by densitometry, four biological repeats, mean ± SEM; ANOVA/Dunnet. **(B)** Levels of WAVE and WASH complexes, based on the amount of the NPF, is illustrated by transparency of the schemes. **(C)** Migration persistence of single cells. Three biological repeats with similar results. Mean ± SEM, n=60 cells. Statistical significance of persistence is analyzed in a non-linear mixed effect model. **(D)** Quantification of ARPC2 enrichment at the surface of endosomes. Three biological repeats with similar results. Mean ± SEM, n=45 cells per condition; Kruskal-Wallis followed by Dunn’s multiple comparison tests. * p<0.05, ** p<0.01, *** p<0.001, **** p<0.0001, ns non-significant.

We then examined whether the crossed effect on the assembly, which is significant, but less pronounced than HSBP1 depletion, is associated with functional defects. Migration persistence that depends on the WAVE complex was significantly decreased upon CCDC53 depletion (Fig.6C, Movie S3). Conversely, endosomal branched actin that depends on the WASH complex was significantly decreased upon BRK1 depletion (Fig.6D). Together these experiments demonstrate that the assembly of a NPF complex does not only depend on its own subunits, but also, surprisingly, on the small subunit of the analogous complex.

## DISCUSSION

We had previously identified HSBP1 as an assembly factor for the WASH complex (Visweshwaran *et al*, 2018). The present study extends the role of HSBP1 to WAVE complex assembly. The fact that HSBP1 promotes the assembly of both WAVE and WASH complexes, is consistent with the fact that these two complexes depend on a similar biochemical reaction: the dissociation of a free heterotrimer of their smallest subunit, BRK1 or CCDC53, to contribute a single molecule to the assembling complex. BRK1 undergoes a transition from a homotrimeric coiled coil precursor to a heterotrimeric coiled coil with its partner subunits, ABI1 and WAVE2, in the final assembled structure of the WAVE complex (Derivery *et al*, 2008; Chen *et al*, 2010).

HSBP1 is concentrated at centrosomes (Visweshwaran *et al*, 2018). The WASH complex, but not the WAVE complex, has been detected associated with centrosomes (Farina *et al*, 2016), suggesting that HSBP1 is likely to assemble the WASH complex at the centrosome (Fokin & Gautreau, 2021). The location of WAVE complex assembly is less clear. WAVE complex subunits are locally translated in membrane protrusions (Willett *et al*, 2013; Mardakheh *et al*, 2015). HSBP1 might promote WAVE complex assembly at membrane protrusions, even if it is not enriched at this subcellular location, the same way it inactivates trimers of the HSF1 transcription factor in the nucleus, where it is also not enriched (Satyal *et al*, 1998). Alternatively, it is possible that neosynthesized proteins encoding subunits of the WAVE complex are transported from membrane protrusions to the centrosome for their assembly. This hypothesis would be consistent with the Nudel adaptor of the dynein microtubule motor being an assembly factor for the WAVE complex that accumulates at centrosomes (Feng *et al*, 2000; Wu *et al*, 2012). The cell biology of multiprotein complex assembly is a challenging question for the future.

WAVE and WASH complex require assembly right after their subunits have been synthesized by ribosomes, but perhaps not only at this time. Indeed, we have recently identified so-called WAVE shell complexes containing a NHS family protein instead of their WAVE subunit in MCF10A cells (Wang *et al*, 2023; Tsydenzhapova *et al*, 2026). NHS and NHSL1 form heterotrimeric coiled coil with ABI1 and WAVE2, like the WAVE NPF subunit. Indirect evidence suggests that embedded NHS family proteins can be exchanged for WAVE subunits. It is possible that such an exchange of subunits corresponds to an assembly event that requires HSBP1. In this study, we systematically measured the levels of the ten subunits forming the two complexes. Clearly, it is the levels of the two NPFs themselves that depend most on HSBP1 levels in depletion and rescue experiments with HSBP1, BRK1 and CCDC53.

Our study reports the existence of various hetero-oligomers containing HSBP1, BRK1 and CCDC53. We managed to reconstitute those from recombinant proteins and to characterize their composition and stoichiometry using native mass spectrometry. The five hetero-oligomers we have identified probably represent a fraction of the many molecular species that these three proteins can form, since their coiled coils appear promiscuous. Native mass spectrometry probably does not allow us to fully explore the space of possible species, since some oligomers may be less stable than others in the gas phase. Importantly, however, we managed, using this approach, to detect the heterotrimer containing one molecule of each and the oligomers containing BRK1 and CCDC53 that can still be formed in the absence of HSBP1. These oligomers were validated in transfected cells using a combination of tags. The important point of these characterizations is that these hetero-oligomers would correspond to the dissociation of BRK1 and CCDC53 homotrimeric free forms, which is a required step for the assembly of their cognate multiprotein complex (Fig.7).

**Figure 7.**
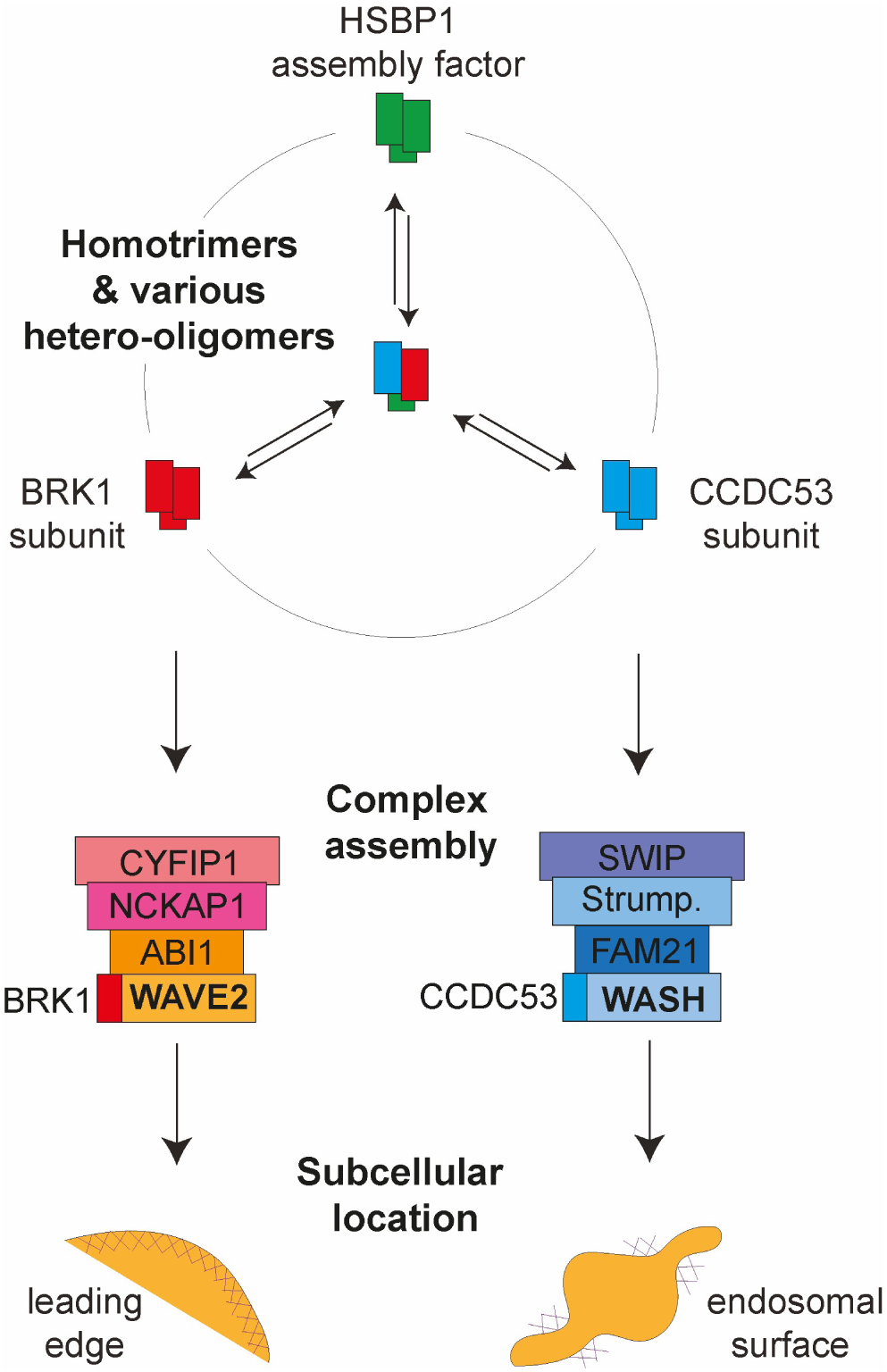
Model of coordination between the assembly of WAVE and WASH complexes through BRK1, CCDC53 and HSBP1 oligomers. The BRK1 precursor promotes WAVE complex assembly and the CCDC53 precursor promotes WASH complex assembly through their dissociation to provide a single subunit to the assembling NPF complex. The HSBP1 assembly factor favors the dissociation of BRK1 and CCDC53 precursors by forming various hetero-oligomers with BRK1 and CCDC53, including a heterotrimer containing one molecule of each.

The function of HSBP1 in assembling both WAVE and WASH complexes is conserved in the amoeba *Dictyostelium* and in human cells. Such a distant evolutionary conservation indicates an essential function. Yet our study also shows that the assembly of these two multiprotein complexes does not absolutely require HSBP1, since WAVE assembly can be rescued by an excess of BRK1, and WASH assembly can be rescued by an excess of CCDC53, in the absence of HSBP1. This conundrum may suggest that the essential function that HSBP1 provides is not precisely the assembly of the two complexes, but rather the coordination between the two assemblies: in knock-outs of HSBP1, the assembly of both complexes concomitantly decreases. But the small subunits BRK1 and CCDC53 also provide coordination between the two assemblies, since in knock-down of BRK1, the WASH assembly - to which BRK1 does not belong - decreases, and reciprocally, in knock-down of CCDC53, the WAVE assembly - to which CCDC53 does not belong - decreases. In other words, the small subunits BRK1 and CCDC53 are subunits of their respective complex, but assembly factors of the other complex to which they do not belong.

In this study, we have revealed the coordinated assembly of the two NPF complexes, but the possible role of such coordination remains elusive. After all, these two molecular machines have diverged to perform unique, non-redundant functions at different subcellular locations, despite the interplay of BRK1 and CCDC53 we report here. One can imagine, however, that the coordination between WAVE and WASH assemblies is important in motile cells, since WAVE-induced cortical branched actin promotes membrane protrusion, whereas WASH-induced endosomal branched actin probably delivers membrane and recycles integrins to the leading edge (Zech *et al*, 2011; Duleh & Welch, 2012; Visweshwaran *et al*, 2018). Both WAVE and WASH complexes play a role in cancer progression and HSBP1 overexpression in tumors might be an efficient way of up-regulating both complexes at once (Visweshwaran *et al*, 2018; Molinie *et al*, 2019). Future studies will probably reveal other instances where the concomitant up-regulation of both NPF complexes is physiologically or pathophysiologically significant. Different cell types can probably tune the level of coordination between NPF complex assembly through HSBP1 expression. To conclude, it seems that nature took advantage of the structural similarity between these two NPF complexes to incorporate the possibility to jointly control their assembly at a common early step.

## METHODS

### Cell Culture

293T cells were from ATCC (CRL-3216). 293 Flip-In T-Rex cells were from Invitrogen. MCF10A cell line was from the collection of breast cell lines organized by Thierry Dubois at Institut Curie, in Paris. 293T and 293 Flip-In T-Rex cells were grown in DMEM medium with 10% FBS and 100 U/mL penicillin/streptomycin. MCF10A cells were grown in DMEM/F12 medium supplemented with 5% horse serum, 20 ng/mL epidermal growth factor, 10 µg/mL insulin, 100 ng/mL cholera toxin, 500 ng/mL hydrocortisone and 100 U/mL penicillin. Media and supplements were from Life Technologies. Cells were incubated at 37 °C in 5% CO2.

*Dictyostelium discoideum* cells were maintained in HL5 medium with glucose (Formedium) in 10 cm Petri dishes at ambient temperature (approx. 22°C).

### Plasmids

The different ORFs encoding subunits WAVE and WASH complexes were shuttled to the custom-made integrative plasmid MXS AAVS1L SA2A Puro bGHpA EF1Flag GFP Blue SV40pA AAVS1R in the place of the Blue cassette using FseI and AscI restriction sites (Molinie *et al*, 2019). All ORFs had previously been cloned between FseI and AscI sites: CYFIP1 (XM_039225), NCKAP1 (AB011159), WAVE2 (AB026542), BRK1 (BC019303) (Gautreau *et al*, 2004) ; ABI1 (FJ380059) in (Derivery *et al*, 2009a) , FAM21 also known as WASHC2C (NM_001330074.2) in (Fokin *et al*, 2021); SWIP also known as KIA1033 and WASHC4 (BC104994.1) in (Ropers *et al*, 2011); WASH also known as WASHC1 (NM_182905) in (Derivery *et al*, 2009b); CCDC53 also known as WASHC3 (NM_016053), Strumpellin also known as WASHC5 (NM_014846) in (Visweshwaran *et al*, 2018).

HSBP1 (BC007515) (Visweshwaran *et al*, 2018) was cloned between FseI and AscI sites and shuttled into the custom-made plasmid MXS EF1Flag Blue SV40pA PGK Blasti BGHpA in the place of the Blue cassette (Visweshwaran *et al*, 2018). BRK1, HSBP1 and CCD53 were similarly shuttled into the pCS2-Flag Blue, pCS2-HA Blue and pCS2-GFP Blue plasmids.

For bacterial expression, HSBP1 was expressed in pET28b (Novagen) either untagged or as His tagged protein; CCDC53 was expressed in pACYC184 (New England Biolabs) either as Flag or HisFlag tagged protein; BRK1 was expressed in pCDF1b (Novagen) either as PC or HisPC tagged protein. All tags were in the N-terminal of proteins.

### Transfection and Isolation of Stable Cell Lines

The calcium phosphate method was used for transient transfection of 293T cells. Cells were analyzed 48 h after transfection.

293 Flp-In T-Rex transfection was performed using Lipofectamine 3000 (Invitrogen). To obtain stable pools expressing Flag-GFP tagged subunits, MXS AAVS1 vectors were co-transfected with two TALEN constructs inducing a double-strand break at the AAVS1 locus (Addgene #59025 and 59026) into 293 Flp-In T-Rex cells, and cells were selected with 0.5 µg/mL puromycin (Invivogen). To obtain Flag-HSBP1 expressing clones, MCF10A transfection was performed using Lipofectamine 3000 (Invitrogen) and transfected cells were selected with 8 µg/mL blasticidin (Invivogen) and single clones were picked up using cloning rings.

### Knock-down and Knock-out

MCF10A knock-down cells were obtained by transfecting 20 nM siRNAs with Lipofectamine RNAiMAX (Invitrogen). siRNAs were Dharmacon ON-TARGET SMART Pools (HSBP1 – L-011294-00-0005; BRK1 – L-017711-01-0005; CCDC53 – L-020900-02-0005). Cells were analyzed 3 days after transfection.

HSBP1 KO MCF10A cell lines were generated with the CRISPR/Cas9 system. gRNA sequences were selected in Genome Browser (https://genome-euro.ucsc.edu) based on the best on/off-targeting ratio. Oligonucleotides encoding the corresponding gRNA sequences were then cloned into the pRG2-GG vector at the BsaI restriction site. The following gRNAs were used:

HSBP1 gRNA1 (5’-GAGCGGACAAACGGAAGTGT-3’),

HSBP1 gRNA2 (5’-GTCACCTCGGTGGTAAGGGA-3’),

ATP1A1 gRNA (5’-GTTCCTCTTCTGTAGCAGCT-3’).

MCF10A cells were cotransfected with the 3 gRNA encoding plasmids in addition to the one encoding Cas9, using Lipofectamine 3000 (ThermoFischer Scientific). After 3 days, cells were selected with 0.5 μM ouabain for 5 days. Individual clones were picked up after 2 weeks of growth using cloning rings. The ouabain control cells were a stable pool of MCF10A cells that were selected to resistant the ouabain treatment.

The HSBP1 KO and random integrant control strains (both Blasticidin-resistant) were derived from the parent Ax2 strain (Dictybase strain ID DBS0235521) and were previously described (Visweshwaran *et al*, 2018).

### Antibodies

The antibodies used were: anti-GFP (11814460001, Roche); anti-Flag (clone M2, Sigma); anti-HA (12CA5, Sigma); anti-VPS35 (B-5, Santa Cruz); anti-ARPC2 (07-227, Millipore); anti-HSBP1 (clone 2C3, Sigma); anti-Strumpellin (C-14, Santa Cruz); anti-NCKAP1 (A305-178A, Bethyl Laboratories); anti-GAPDH (AM4300, Thermo Fisher Scientific); anti-α-tubulin (T9026, Sigma). Home-made SWIP, WASH, CCDC53, FAM21, CYFIP1, ABI1, WAVE2 antibodies and BRK1 antibody were previously described (Gautreau *et al*, 2004; Derivery *et al*, 2008, 2009b). Home-made *Dictyostelium* antibodies were polyclonal antibodies anti-Scar developed in sheep (Pollitt *et al*, 2006) or in rabbit (#9618) (Ura *et al*, 2012).

### Western Blots

Cells were lysed in RIPA buffer (HEPES 50 mM, EDTA 10 mM, SDS 0.1%, NP-40 1%, DOC 0.5%, NaCl 15 mM, pH7.4) supplemented with protease inhibitors (Roche). Lysates were clarified by centrifugation at 16,000 xg for 15 min and subjected to SDS–PAGE using NuPAGE 4-12% Bis-Tris gels (Life Technologies). An optimized protocol of Western Blot was specifically used (Visweshwaran *et al*, 2018).

Pieces of nitrocellulose membranes were cut and incubated with different primary antibodies, HRP conjugated secondary antibodies (Sigma) and developed with SuperSignal™ West Femto Substrate (Thermo Fisher Scientific) and ChemiDoc imaging system (Bio-Rad). For the characterization of 293 Flip-In stable cell lines expressing Flag-GFP tagged subunits of WAVE and WASH complexes, the secondary antibodies used were conjugated with alkaline phosphatase (Promega), and nitrocellulose membranes were developed using NBT/BCIP as substrates (Promega). For *Dictyostelium* Western blots, membranes were developed with either secondary rabbit anti-sheep (cross-adsorbed dylight 800 IgG (H+L); ThermoFisher Scientific, SA5-10060) or goat anti-rabbit antibodies (ThermoFisher Scientific, SA5-10036) antibody and imaged on a LiCor Odyssey CLX. Bands were quantitated in the LiCor ImageStudio software. Blots were re-probed using Streptavidin-AlexaFluor680 (ThermoFisher Scientific, S21378) to detect the biotinylated MCCC1 loading control (Davidson *et al*, 2013), used for normalization.

### Immunoprecipitation and Tandem Affinity Purification

Cells were lysed in XB-NP40 buffer (50 mM HEPES, 50 mM KCl, 1% NP-40, 10 mM EDTA, pH 7.7) supplemented with protease inhibitors (5056489001, Sigma) at 4 °C for 30 min. The lysates were clarified by centrifugation at 16,000 xg for 15 min. For GFP immunoprecipitation, clarified cell extracts were incubated with GFP-trap beads (Chromotek) at 4 °C for 1 h. The GFP-trap beads were washed with the XB-NP40 buffer. Beads were subjected to SDS–PAGE for Western blot.

For tandem affinity purification clarified cell extracts were incubated with FLAG-M2 beads (Sigma) at 4 °C for 4 h. FLAG-M2 beads were washed with the XB-NP40 buffer and eluted with 0.5 mg/mL 3xFLAG peptide (F4799, Sigma) in XB (50 mM HEPES, 50 mM KCl, 10 mM EDTA, pH 7.7) overnight at 4 °C. FLAG elutions were then incubated with GFP-trap beads (Chromotek) at 4 °C for 1 h. The GFP-trap beads were washed with the XB-NP40 buffer. 20% of the beads were subjected to SDS–PAGE and colloidal Coomassie staining followed by silver staining (SilverQuest Silver Staining Kit, Thermo Fisher Scientific). The remaining 80% were analyzed by mass spectrometry.

### Immunofluorescence

Cells were seeded on glass coverslips previously coated with 20 µg/mL Fibronectin (Sigma) for 1 h at 37°C. Cells were fixed in 3.2% PFA then permeabilized with 0.5 % Triton X-100, blocked in 2% BSA and incubated with antibodies (1-5 µg/mL for the primary, 5 µg/mL for the secondary). Cells were fixed in 4% PFA then permeabilized with 0.5 % Triton X-100, blocked in 2% BSA and incubated with primary VPS35 and ARPC2 primary antibodies (anti-VPS35 and then with Alexa Fluor 488 or 555 conjugated secondary antibodies and DAPI (Life Technologies). Images were acquired using a SP8ST-WS confocal microscope (Leica) equipped with a HC PL APO 63x/1.40 oil immersion objective, a white light laser, HyD and PMT detectors. Image analysis was performed using the ImageJ FIJI software. To quantify the endosomal enrichment of ARPC2 staining, images were thresholded in the VPS35 channel using the Otsu algorithm to identify VPS35-positive endosomal structures and generate regions of interest (ROIs). The mean ARPC2 fluorescence intensity was measured within the VPS35-positive ROIs. As a local cytoplasmic reference, the ROIs (of same size and shape) were repositioned to VPS35-negative cytoplasmic regions within the same cell, and the mean ARPC2 fluorescence intensity was measured. Endosomal enrichment of ARPC2 was calculated as the ratio of the mean ARPC2 intensity within VPS35-positive endosomal ROIs to that measured in cytoplasmic ROIs.

### Migration and Videomicroscopy

Cell migration assays with MCF10A cells were performed in μ-Slide eight-well dishes (#80826, Ibidi), coated with 20 μg/ml Fibronectin (Sigma). Videomicroscopy was performed 24 h after seeding using an inverted Axio Observer microscope (Zeiss) equipped with a Pecon Zeiss incubator XL multi S1 RED LS (Heating Unit XL S, Temperature module, CO2 module, Heating Insert PS and CO2 cover), a definite focus module and a Hamamatsu camera C10600 Orca-R2. Images were acquired every 10 min for 24 h with a ×10 objective. Individual cells were tracked using the Manual Tracking plug-in in the ImageJ software. The DiPer software was used to represent cell trajectories (Plot_at_origin.txt) and analyze the single cell migration parameters: MSD (MSD.txt) and migration persistence (Autocorrel.txt) (Gorelik & Gautreau, 2014). Persistence represents the direction autocorrelation, characterizing the orientation of two displacement vectors, i.e. the cosine of the angle they form, as a function of the time interval that separates these displacement vectors.

Chemotaxis assays with *D. discoideum* were performed using a previously described under-agarose assay (Tweedy *et al*, 2016). 6-well tissue culture dishes were pre-coated with 2% bovine serum albumin solution for 5min, which was then removed. 0.4% molecular biology grade agarose (Melford Laboratories, A20090) was prepared in KK2 buffer (lacking salts) by boiling until dissolved. Once sufficiently cooled, folic acid (Merck Life Sciences F7876) was added to a final concentration of 4 µM (from a 10mM stock), 2 ml dispensed into each well, and allowed to set for 1 h. A channel (approx. width 3mm) was then cut vertically into each well. Cells were harvested from Petri dishes by gentle aspiration, centrifuged at 300 xg for 3 min, washed twice in KK2, and resuspended at 2x10^6^ cells/ml. Approx. 80 µl of cell suspension was carefully pipetted into each channel. Dishes were incubated in a humid box at ambient temperature for 3-4 h to allow cells to begin chemotaxis and move sufficiently far from the well. For timelapse imaging an inverted Nikon TiE microscope equipped with moving stage (ASI) was used, acquisitions being controlled by Metamorph software. A 10x/0.3NA Plan Fluor Ph1 DLL objective (WD 16mm) was used, images taken with a Teledyne Photometrics Prime camera (native pixel size 6.5µm) using 2x pixel binning (image pixel size 1.3µm). Frame intervals of 60 sec were used. Images were opened in FIJI (Schindelin *et al*, 2012), and cells tracked using a custom plugin.

### Statistical Analysis

Cell speed was averaged for each cell and the averages for each cell were compared across conditions by the Kruskal-Wallis test followed by Dunn’s post-hoc test with Holm correction for multiple comparisons in MCF10A cells and using Student’s t-test for *Dictyostelium*. This analysis was performed using GraphPad Prism.

Directional persistence was quantified as the direction autocorrelation of each cell’s trajectory (Gorelik & Gautreau, 2014). For each condition, the decay of the autocorrelation was fitted to A(t) = exp(-t/τ) using a nonlinear mixed-effects model implemented with the nlme package (Pinheiro & Bates, 2000) in R (R Foundation for Statistical Computing, Vienna, Austria), with the decay rate parameterized on a log scale. Genetic condition was entered as fixed effects on log(1/τ), and a random effect on log(1/τ) was included to account for repeated measurements within cells. Conditions were compared pairwise by fitting, for each pair, a model in which the decay rate (1/τ) was allowed to differ between the two conditions; the p-value tests the null hypothesis of equal decay rate (equal persistence time τ). Visual inspection of residual plots and of the distribution of random effects did not reveal any obvious deviations from homoscedasticity or normality. The resulting p-values were adjusted for multiple comparisons by the Benjamini-Hochberg (FDR). The analysis was made in Microsoft Excel, Python and R.

Differences were considered significant at confidence levels greater than 95% (p < 0.05). Four levels of statistical significance were distinguished: p < 0.05; p < 0.01; p < 0.001; p <0.0001.

No statistical method was used to predetermine sample size. No outlier points were excluded from analyses. The experiments were not randomized. The investigators were not blinded to allocation during experiments and outcome assessment.

## Acknowledgments

This work was supported by grants from Agence Nationale de la Recherche (ANR-22-CE13-0041 and ANR-24-CE44-4957 to AMG), Institut National du Cancer (INCA_16712 to AMG) and Fondation ARC pour la Recherche sur le Cancer (ARC PJA 2023 070006853 to AMG). This work benefited from the support of the “*Precision Biomedicine of Cancer”* led by l’X – Ecole polytechnique and the Fondation de l’Ecole polytechnique, sponsored by “Science et Technologie”, an association of the Servier Group. JCR, LKF and MD would like to acknowledge Bruker for providing access to the technical support for the timsOmni mass spectrometry platform and Dr. Philip Compton (Integrated Protein Technologies) for his support on the SampleStream platform.

## Author Contributions

**Nikita M. Novikov**: Investigation, Writing - Original Draft, Visualization. **Christine Lazennec-Schurdevin, Peter A. Thomason, Iman Haddad, Lucile Kogey-Fuchs, Magalie Duchateau, Daria Bondar, Nicolas Dugave, Sergio Lilla**: Investigation. **Artem I. Fokin:** Investigation, Supervision. **Julia Chamot-Rooke, Joëlle Vinh, Robert H. Insall, Raphaël Guérois, Yves Mechulam, Emmanuelle Schmitt:** Supervision. **Raphaël Guérois:** Conceptualization, Funding acquisition. **Alexis M. Gautreau**: Conceptualization, Data Curation, Funding acquisition, Supervision, Project administration, Writing - Review & Editing.

NMN performed most experiments and analyses. DB and ND were rotating students from Ecole Polytechnique, who generated and characterized stable 293 cell lines under the supervision of AIF. CLS performed the biochemical reconstitution of the HSBP1-CCDC53-BRK1 heterotrimer. PAT and RHI performed the experiments with the amoeba *Dictyostelium*. SL performed MS with *Dictyostelium* samples. IH and JV performed mass spectrometry. CLS, YM and ES contributed the reconstitution of recombinant complexes. LKF, MD and JCR performed native mass spectrometry with reconstituted complexes. RG performed structural modeling. RHI, JCR, JV, YM, ES and AMG supervised the work in their respective groups. AMG coordinated the collaborative work and wrote the manuscript.

## Supporting Information

Supplementary information is available.

## Data availability

Source data are provided with this paper. Raw files of mass spectrometry have been deposited in PRIDE with access for the reviewers.

WAVE data: PXD070545 Token rbdy4rHkgGLL WASH data: PXD070960 Token 7vydVrVmYPpK *Dictyostelium* data: PXD083888 Token bprM3EXgvgb9

Native mass spectrometry of reconstituted complexes: PXD083996 Token Ff0R6CV41V8e

## Abbreviations used

au: arbitrary unit
bp: base pairs
cl.: clone par. parental
EV: Empty Vector
LFQ: Label Free Quantification
MSD: Means Square Displacement
NPF: Nucleation Promoting Factor
na: not applicable
nd: not determined
ns: non significant
Strump.: Strumpellin
TAP: Tandem Affinity Purification

